# Interferon-induced Protein-44 and Interferon-induced Protein 44-like restrict replication of Respiratory Syncytial Virus

**DOI:** 10.1101/2020.02.24.963900

**Authors:** D. C. Busse, D. Habgood-Coote, S. Clare, C. Brandt, I. Bassano, M. Kaforou, J. Herberg, M. Levin, Jean-Francois Eleouet, P. Kellam, J. S. Tregoning

## Abstract

Cellular intrinsic immunity, mediated by the expression of an array of interferon-stimulated antiviral genes, is a vital part of host defence. We have previously used a bioinformatic screen to identify two interferon stimulated genes (ISG) with poorly characterised function, Interferon-induced protein 44 (IFI44) and interferon-induced protein 44-like (IFI44L), as potentially being important in Respiratory Syncytial Virus (RSV) infection. Using overexpression systems, CRISPR-Cas9-mediated knockout, and a knockout mouse model we investigated the antiviral capability of these genes in the control of RSV replication. Over-expression of IFI44 or IFI44L was sufficient to restrict RSV infection at an early time post infection. Knocking out these genes in mammalian airway epithelial cells increased levels of infection. Both genes express antiproliferative factors that have no effect on RSV attachment but reduce RSV replication in a minigenome assay. The loss of *Ifi44* was associated with a more severe infection phenotype *in vivo*. These studies demonstrate a function for IFI44 and IFI44L in controlling RSV infection.

**Importance:** RSV infects all children under two years of age, but only a subset of children get severe disease. We hypothesize that susceptibility to severe RSV necessitating hospitalization in children without pre-defined risk factors is in part mediated at the anti-viral gene level. But there is a large array of anti-viral genes, particularly in the ISG family about which the mechanism is poorly understood. Having observed significantly lower levels of IFI44 and IFI44L gene expression in hospitalized children with a confirmed diagnosis of RSV, we dissected the function of these two genes. Through a range of over-expression and knockout studies we show that the genes are anti-viral and anti-proliferative. This study is important because IFI44 and IFI44L are upregulated after a wide range of viral infections and IFI44L can serve as a diagnostic bio-marker of viral infection.

## Introduction

Respiratory syncytial virus (RSV) is a major global cause of morbidity in young children and the elderly, representing a significant burden on healthcare infrastructure (1). The majority of RSV infections in susceptible populations are self-limiting, however some 2% of infected infants develop a severe infection and require hospitalisation. The risk factors behind this development of severe RSV disease have yet to be fully elucidated and some 75% of hospitalised infants present with no known risk factor (2, 3). This suggests that there is a genetic element to susceptibility to symptomatic infection and since ISGs are vital in early viral control, they are a likely candidate. A number of ISG have been demonstrated to inhibit RSV including IFITM proteins (4, 5) TDRD7 (6), and 2′-5′ oligoadenylate synthetase (7).

A vital component of the innate host response to viral infection is the intracellular amplification of an array of antiviral proteins in response to type I interferon (IFN). The majority of these inducible proteins, encoded by IFN-stimulated genes (ISGs), have no defined function and have only been poorly characterised in terms of their antiviral tropism. Understanding how these genes reduce viral infection gives insight into the viral life cycle and may open up novel therapeutic routes. We have previously performed a bioinformatic screen of ISG expressed after RSV infection, which prioritised ISG of interest for further study (8).

Two ISGs of interest identified in our previous work are *IFI44* and *IFI44L*, which are found adjacently on chromosome one. *IFI44L* is a larger gene of 26 kilobases (kb) compared to the 14 kb of *IFI44*, but both genes encode similar sized proteins translated from a transcript produced from nine exons. IFI44 is made up of 444 amino acids whereas IFI44L has 452 residues: the two proteins share 45% amino acid identity. *IFI44*, previously known as *MTAP44*, was first identified in the context of Hepatitis C virus infection (9, 10). Overexpression of IFI44 has been shown to restrict Bunyamwera virus (11), and HIV-1 (12) infection *in vitro*. IFI44 was initially described as a cytoplasmic protein, however two studies have observed that low levels of IFI44 can be found in the nucleus (12, 13). Hallen *et al.* reported that overexpression of IFI44 was able to reduce proliferation of two melanoma cell lines independently of IFN-I (13). The anti-proliferative mechanism of IFI44 remains unexplored.

Even less is known about the tropism and function of IFI44L. IFI44L has been shown to have a moderate impact on Hepatitis C virus infection (14). Interestingly, *IFI44L* expression has also been associated with several autoimmune disorders (15–17), cancer (18, 19), and humoral responses to vaccination (20). These seemingly disparate contexts suggest that *IFI44L* may be a biomarker of IFN responses independent of the type of stimulus. Interestingly, IFI44L expression is sufficient to distinguish viral from bacterial infection (21). Like IFI44, IFI44L has antiproliferative activity, associated with increased activation of Met/Src signalling (18).

Using both overexpressing and knockout cell lines, we demonstrated that IFI44 and IFI44L are antiproliferative factors that can independently restrict RSV infection. We report that this ability to restrict infection involves the reduction of viral genome transcription or replication but was not dependent upon a predicted guanosine-5’-triphosphate (GTP)-binding region present in either protein. We demonstrate for the first time that the loss of IFI44 expression in a mouse model of infection is associated with more severe RSV disease.

## Methods

### Clinical cohort

*IFI44* and *IFI44L* expression was analysed in a published clinical cohort of febrile infants with either moderate or severe RSV infection [13]. The microarray gene expression dataset (GSE72810) was retrieved from the National Institutes of Health Gene Expression Omnibus database (22) using the GEOquery package (23) in R (24). Normalisation was performed using robust spline normalisation (RSN) from the lumi package (25) followed by a log transformation. Patients with suspected or confirmed bacterial infection were removed (n = 4). Cohort demographics are described in the associated figures. Prior to differential expression probes were removed if the expression was not above 6 in at least 4 samples. Differential expression was performed using Limma and expression values were normalised using robust spline normalisation and a log transformation, plotted p-values are not adjusted for multiple testing (values in tables are Benjamini-Hochberg corrected). Interferon stimulated genes were downloaded from KEGG.

### Cell culture and viruses

HEp-2 (from P. Openshaw, Imperial College London), A549 (ATCC CCL-185), and HEK293T/17 (ATCC CRL-11268) cells were maintained in Dulbecco’s modified eagle medium supplemented with 10% v/v foetal calf serum, 1% v/v penicillin/streptomycin, and 1% v/v L-glutamine. RSV strain A2 (from P. Openshaw, Imperial College London) and rgRSV (26) were passaged in HEp-2 cells before quantification of viral titre by plaque assay. VSV-G pseudotyped Lentiviral particles were produced by triple transfection in HEK293T/17 cells using Lipofectamine 3000 (Thermofisher). Lentiviral vectors pTRIP-FLUC-tagRFP, pTRIP-IFI44-tagRFP, and pTRIP-IFI44L-tagRFP were a kind gift from M. Dorner (Imperial College London). To generate mutant proteins these vectors were altered by using QuickChange® XL site-directed mutagenesis (Agilent Technologies) according to the manufacturer’s instruction. Lentivirus was harvested 24-52 hours after transfection and concentration of transducing units (TU) determined by flow cytometry. For transduction, 5×10^4^ cells were seeded into each well of a 24-well plate. After 24 hours cells were transduced with 2×10^5^ TU ml^−1^ lentivirus containing supernatant. Stably transduced clonal populations were recovered following fluorescence-activated cell sorting (FACS). RFP expression was monitored over three weeks and expression of IFI44 or IFI44L confirmed by quantitative PCR (qPCR) or Western blotting.

### Quantitative PCR

For analysis of *in vitro* samples, cells were lysed in RLT buffer and RNA extracted using Qiagen RNeasy kit according to the manufacturer’s instructions (Qiagen). RSV viral load *in vivo* was assessed by extracting RNA from frozen lung tissue using Trizol extraction after disruption in a TissueLyzer (Qiagen). Complementary DNA (cDNA) was reverse transcribed from RNA extracts using GoScript reverse transcriptase with random primers according to the manufacturer’s instructions (Promega). qPCR reactions were carried out on a Stratagene Mx3005p thermal cycler (Agilent Technologies). RSV viral load was quantified by amplification of the RSV L gene using 900 nM forward primer (5’-GAACTCAGTGTAGGTAGAATGTTTGCA-3’), 300 nM reverse primer (5’-TTCAGCTATCATTTTCTCTGCCAAT-3’), and 100 nM probe (5’-FAM-TTTGAACCTGTCTGAACAT-TAMRA-3’) in Taqman™ Universal master mix, no AmpErase™ UNG (Thermofisher). Absolute copy number was calculated by comparison to a plasmid standard. mRNA was amplified with SYBRselect master mix (Thermofisher) according to the manufacturer’s instructions. The following primers were used at a final concentration of 250 nM: hIFI44 (forward 5’-TGGTACATGTGGCTTTGCTC-3’, reverse 5’-CCACCGAGATGTCAGAAAGAG-3’), hIFI44L (forward 5’-AAGTGGATGATTGCAGTGAG-3’, reverse 5’-CTCAATTGCACCAGTTTCCT-3’), hGAPDH (forward 5’-GGACCTGACCTGCCGTCTAG-3’, reverse 5’-TAGCCCAGGATGCCCTTGAG-3’), m*Ifi44* (forward 5’-AACTGACTGCTCGCAATAATGT-3’, reverse 5’-GTAACACAGCAATGCCTCTTGT-3’), m*Ifi44l* (forward 5’-AGTGACAGCCAGATTGACATG-3’, reverse 5’-CATTGTGGATCCCTGAAGAGAA-3’), and m*Gapdh* (forward 5’-AGGTCGGTGTGAACGGATTTG-3’, reverse 5’-TGTAGACCATGTAGTTGAGGTCA-3’). Fold-change in target gene in treated samples was calculated using the ΔΔCt method (Ct = cycle threshold) and normalised to a reference transcript (27).

### Western blotting

Cells were lysed using RIPA buffer (Sigma) containing 1 x cOmplete™ Ultra protease inhibitor cocktail (Sigma). Proteins were separated by SDS-PAGE using 4-20% pre-cast Mini-Protean TGX gels (Bio-rad) before transfer onto a nitrocellulose membrane using the Trans-Blot turbo transfer system (Bio-rad). Membranes were blocked for 1 hour at room temperature (RT) with 5% milk in PBS with 0.1% Tween 20. Membranes were probed using the following primary antibodies for 16 hours at 4°C: IFI44 (ThermoFisher, PA5-65370) and IFI44L (VWR, ARP46166), β-actin (Abcam, ab8227). Membranes were washed and probed with anti-IgG HRP-conjugated antibodies (Dako) prior to chemiluminescent detection.

### Flow cytometry

Analysis was performed on a Becton Dickinson Fortessa LSR using a 561 nm laser and 582/15 band pass filter to detect tag-RFP positive cells, a 499 nm laser and 530/30 band pass filter to detect GFP positive cells. Acquisition was set to record 10^4^ events followed by doublet gating and analysis with FlowJo V10.

### RSV Plaque assay

RSV titre in cell-free supernatant was quantified by immunoplaque assay using biotinylated goat anti-RSV polyclonal antibody (Abcam). HEp-2 cells were infected with dilutions of RSV-containing supernatant for 24 hours. Cells were fixed in methanol containing 2% hydrogen peroxide for 20 minutes at RT. Cells were washed with 1% bovine serum albumin (BSA) PBS prior to addition of anti-RSV antibody for 1 hour. Plaques were then visualised by incubating the cells with ExtrAvidin peroxidase followed by 3 amino-ethylcarbazole substrate (Sigma).

### CRISPR knockout generation

Guide RNA (gRNA) sequences targeting human *IFI44* (gRNA1: CAA TAC GAA TTC T; gRNA2: GAA AGA AGG CGG CCT GTG C) and *IFI44L* (gRNA1: TAA CCT AGA CGA CAT AAA G; gRNA2: GTG ACT GGC CAA GCC GTA G) were synthesised according to Sanjana et al (28) and cloned into pSpCas9(BB)-2A-GFP (Addgene, #48138) following *Bbs*I digestion. Insertion of gRNA sequences validated by sequencing (Eurofins Genomics). A549 cells were transfected and sorted by FACS after 48 hours. Clonal knockouts were validated by PCR amplification of the targeted region, agarose gel electrophoresis, Western blotting, and sequencing. Clustal Omega was used for multiple sequence alignment (29).

### Proliferation assays

Viable cell numbers were quantified either by Trypan blue exclusion or by the production of formazan product (OD^490^) 2 hours after addition of CellTiter 96^®^ Aqueous One Solution assay reagent (Promega) according to the manufacturer’s instructions. Different cell lines were seeded at equal densities and viable cell number quantified 6-48 hours later. Cell division was assessed by staining cells with 5 μM CellTrace Violet reagent (Thermofisher) and flow cytometric analysis (10^4^ single cell events, 405 nm laser with a 450/50 band pass filter) after 72 hours. Prior to analysis cells were harvested and fixed for 20 minutes at RT in 4% paraformaldehyde.

### RSV cold-bind assay

Stably transduced cell lines were seeded at equal densities 24 hours prior to infection. Each cell line was counted prior to infection to ensure inoculums were normalised across different cell lines. Cells were equilibrated to 4°C for 30 minutes before media was removed and infected with RSV A2 in a minimal volume of serum-free DMEM for 90 minutes at 4°C. Cells were then washed 3 x with ice-cold 1x PBS and lysed in RLT buffer (Qiagen) with 1/100 β-mercaptoethanol (Sigma) for quantification of RSV L gene RNA.

### RSV minigenome assay

The RSV Minigenome and plasmids expressing RSV L, N, P, and M2-1 proteins were described previously (30). pGEM3-Gaussia/Firefly encodes a sub-genomic RSV replicon. From the 3’ end: A2 leader sequence (Le), Gaussia luciferase open-reading frame (ORF) with an NS1 gene start (GS) and M gene end (GE) sequence, Firefly luciferase ORF with SH GS and GE sequences, A2 trailer region (Tr). HEK293T/17 cells and stably transduced 293T/17 cell lines were seeded 24 hours prior to transfection at 90% confluency in 24-well plates. The cells were transfected using Lipofectamine 3000 with a DNA mixture of 0.25 μg pGEM3-Gaussia/Firefly minigenome, 0.125 μg pCITE-L, 0.25 μg pCITE-P, 0.06 μg pCITE-M2-1, 0.25 μg pCITE-N, 0.12 μg pSV-β-Gal (Promega), and 0.25 μg pCAGGS-T7 (Addgene #65974). Negative controls were transfected with the DNA mix with pCITE-L replaced by pcDNA3.1. Cells were lysed in 1 x passive lysis buffer (Promega) after 24 hours. Firefly luciferase activities were measured in 10 μl of lysate using 50 μl luciferase assay substrate (Promega). To normalise transfection efficiencies β-Galactosidase levels were measured using the β-Galactosidase Enzyme Assay System (Promega). 20 μl of lysate was diluted 1:1 in 1x reporter lysis buffer before addition of 40 μl 2x assay buffer. Following incubation at 37°C for 1 hour 150 μl 1 M Na_2_CO_3_ was added and absorbance measured at 420 nm.

### Mouse infection

Background-, sex-, and age-matched, >95% C57BL/6N or BALB/c, wild type and *Ifi44*^*tm1b(komp)Wtsi*^ (*Ifi44*^*−/−*^) mice (31) (Wellcome Trust Sanger Institute) were supplied with food and water *ad libitum* and monitored daily. Mice were infected intranasally (i.n.) with 1-4 × 10^5^ plaque-forming units (PFU) of RSV A2 in 100 μL under isoflurane anaesthesia.

### Enzyme-linked immunosorbent assay

Bronchoalveolar lavage fluid (BALF) was collected by inflating the lungs with PBS. Supernatant was collected after centrifugation and assayed. For lung homogenate, lung tissue was homogenised through a 100 μm cell strainer (Falcon) and supernatant collected after centrifugation and ACK lysis. Cytokines in lung homogenate and BALF were quantified using Duoset ELISAs according to the manufacturer’s instructions (R&D systems).

### Luminex^®^ multiplex ELISA

Lung homogenate supernatant was subjected to a magnetic Luminex^®^ assay using a premixed multi-analyte kit: CXCL10, CCL3, CXCL2, CXCL1, IL-5, CCL2, IL-1α, and CCL5 (R&D systems). Samples and diluted microparticles were combined according to the manufacturer’s instructions and incubated for 2 hours at RT (800 rpm). A magnet was applied to the bottom of the plate and wells were washed x 3 for 1 minute each, before addition of a Biotin Antibody cocktail (1h RT, 800 rpm). The previous wash was repeated, and Streptavidin-PE added to each well (30m RT, 800 rpm). The wash was repeated and microparticles resuspended in wash buffer for analysis on a Bio-Plex^®^100 Luminex machine (Bio-rad).

### Statistical analysis

Analysis performed in Prism 8 as described in figure legends (GraphPad Software).

### Ethics

All animal experiments were maintained in accordance with UK Home Office regulations, the UK Animals (Scientific Procedures) Act 1986 and reviewed by an Animal Welfare and Ethical Review Body. The work was done under PPL P4EE85DED. Clinical data presented was collected in a previous study [13]. Written informed consent was obtained from parents or guardians using locally approved research ethics committee permissions (St Mary’s Research Ethics Committee (REC 09/H0712/58 and EC3263); Ethical Committee of Clinical Investigation of Galicia (CEIC ref 2010/015); UCSD Human Research Protection Program No. 140220; and Academic Medical Centre, University of Amsterdam (NL41846.018.12 and NL34230.018.10).

## Results

### IFI44 and IFI44L expression in human infants with severe RSV infection

We have previously identified a number of ISGs that are consistently upregulated in response to RSV (8), of which IFI44 and IFI44L featured prominently. We focussed on these two genes as relatively little was known of their phenotype. Our hypothesis was that at an individual gene level, there should be a significant difference in gene expression RSV infected children with severe disease. To test this, we examined gene expression levels of both *IFI44* and *IFI44L* in data previously generated by microarray on RNA extracted from whole blood derived PBMCs, collected from children with confirmed RSV infection. We compared those who required paediatric intensive care unit (PICU) admission with those who were admitted to a general hospital ward (General). When investigated as individual genes in the dataset, the expression of both IFI44 (P = 0.0082, Fig. 1b) and IFI44L (P = 0.0248, Fig. 1c) was significantly lower in those patients admitted to the paediatric intensive care unit (PICU). IFI44 expression correlated with IFI44L expression across both moderate and severe RSV patients (p<0.001, r^2^=0.74, Fig. 1d). It should be noted that, when investigated in the context of global gene expression data, the differences between general hospital and intensive care admission were not significant, though IFI44L did have a greater than 2log fold change (Fig. 1e). This observed difference supported the rationale for looking at the function of IFI44 and IFI44L in more detail.

**Figure 1.**
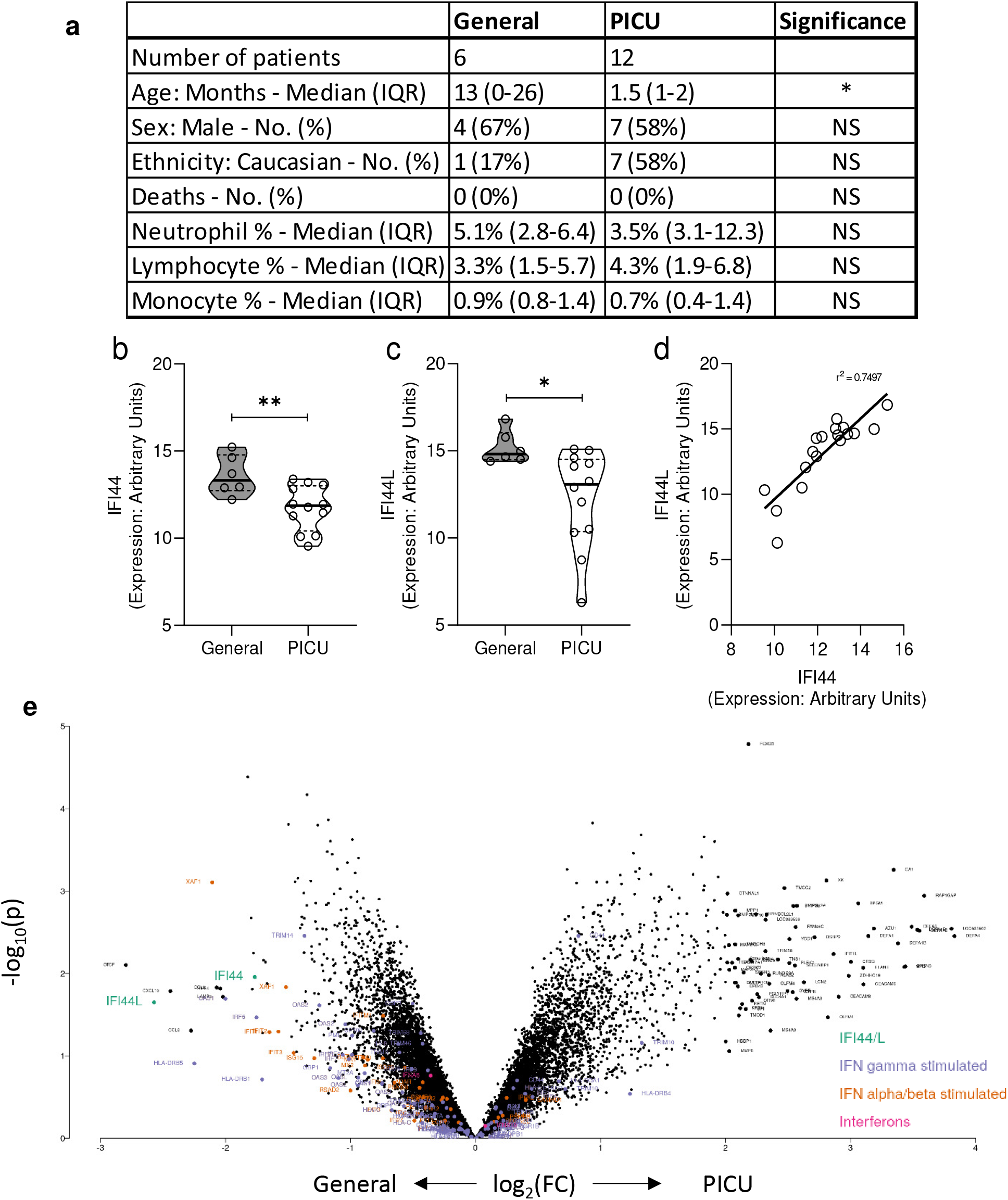
IFI44 and IFI44L mRNA levels are reduced in the blood of infants with severe RSV infection. **(a)** Demographic and clinical features of patient groups. Patients were febrile children with immunofluorescence-confirmed RSV infection. Patients with suspected or confirmed bacterial co-infection were excluded (n = 4). **(b)** IFI44 or **(c)** IFI44L RNA expression levels, measured by in blood PBMC by microarray in either patients admitted to a general ward with mild RSV illness or admitted to a paediatric intensive care unit (PICU) at the same hospital. * P < 0.05, ** P < 0.01. Significance by unpaired t-test. **(d)** Pearson correlation analysis of expression of IFI44 and IFI44L. P < 0.001. **(e)** Volcano plot of fold change in gene expression by microarray between general hospital and intensive care admission.

### *IFI44* and *IFI44L* are upregulated early in response to IFN-I and RSV

Human lung epithelial A549 cells treated with recombinant IFNα2a robustly upregulated expression of *IFI44* and *IFI44L* mRNA within 2 and 6 hours respectively (P < 0.01, Fig. 2a). Induction following RSV infection was slower than IFN, with *IFI44* and *IFI44L* upregulated within 6 and 12 hours (P < 0.05, Fig. 2b). Expression remained upregulated for at least 48 hours following treatment or infection. IFI44 protein was increased following 24-48 hours of IFNα2a stimulation (Fig. 2c) and undetectable in unstimulated cells. IFI44L protein was detectable in unstimulated cells, it did not appear to be induced by IFN treatment relative to the control untreated cells (Fig. 2c). After recombinant IFN treatment expression of both genes plateaued after 6 hours but expression continued to increase during the first 48 hours of RSV infection, in parallel with viral RNA levels (Fig. 2d). When cells were pre-treated with IFNα2a before infection, there was a significant reduction in viral RNA levels (Fig. 2e)

**Figure 2.**
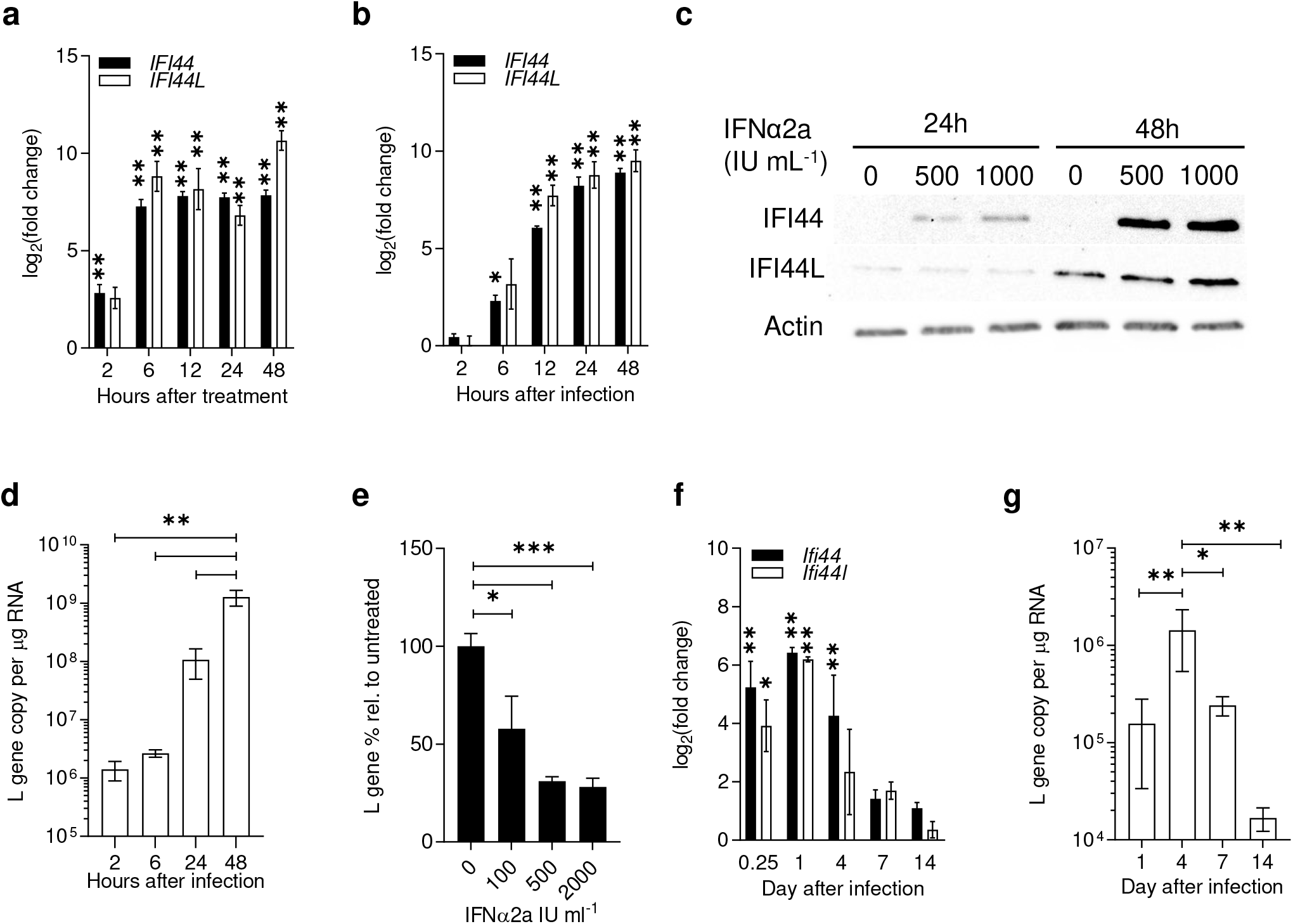
*IFI44* and *IFI44L* are IFN-I responsive genes upregulated during RSV infection. **(a)** *IFI44* and *IFI44L* mRNA expression in A549 cells treated with 500 IU mL^−1^ IFNα2a or **(b)** infected with RSV A2 (m.o.i. 0.1) for 2-24 hours. N≥3. **(c)** IFI44 and IFI44L protein levels in A549 cells treated with 0-1000 IU mL^−1^ IFNα2a for 24 or 48 hours. **(d)** RSV L gene copies in RSV infected A549 cells as for **b**. N≥3. **(e)** A549 cells treated with IFNα2a for 16 hours prior to infection with RSV A2 (m.o.i. 0.1) for 24 hours. Viral RNA relative to untreated controls. N=3. **(f)** 8-10 week old BALB/c mice were infected intranasally with 2 × 10^5^ pfu RSV A2. Change in expression of *Ifi44* and *Ifi44l* mRNA in infected mice relative to mice given PBS intranasally at the time of infection. **(g)** Total L gene copy number per μg RNA from whole lung tissue. N≥4 animals per group at each time point. Data are presented as the mean +/− SEM. Significance relative to untreated controls (**a, b, e**), PBS-treated (**f**), or noted groups (**d**, **g**) assessed by ANOVA. * P < 0.05, ** P < 0.01, *** P < 0.001, **** P < 0.0001.

Similarly, the expression of both *Ifi44* and *Ifi44l* RNA was significantly upregulated rapidly after intranasal infection of BALB/c mice, detectable from 6 hours (P < 0.05, Fig. 2f). Gene expression peaked after 24 hours and returned towards baseline levels by day 14 in parallel with levels of viral RNA that also reduced as the infection was cleared (Fig. 2g).

### Overexpression of IFI44 or IFI44L restricts RSV infection

In a previous screen to identify ISGs that impact RSV infection McDonald *et al.* showed that transient overexpression of human IFI44 or IFI44L by lentiviral transduction reduced the percentage of RSV infected cells (8). Here, we generated stably transduced clonal cell lines by lentiviral transduction, followed by fluorescence-activated cell sorting (FACS) selection. Following expansion, clonal populations were selected that expressed TagRFP after three weeks in culture following sorting. Individual cell lines were then selected based on detectable expression of either *IFI44* or *IFI44L* by qPCR (Fig. 3a). *MX1* expression was assessed in each stable cell line to confirm the specificity of overexpression to either *IFI44* or *IFI44L*. There were no significant differences in expression of *MX1* in either cell line following IFN-I stimulation suggesting that neither IFI44 or IFI44L are regulators of the IFN response.

**Figure 3.**
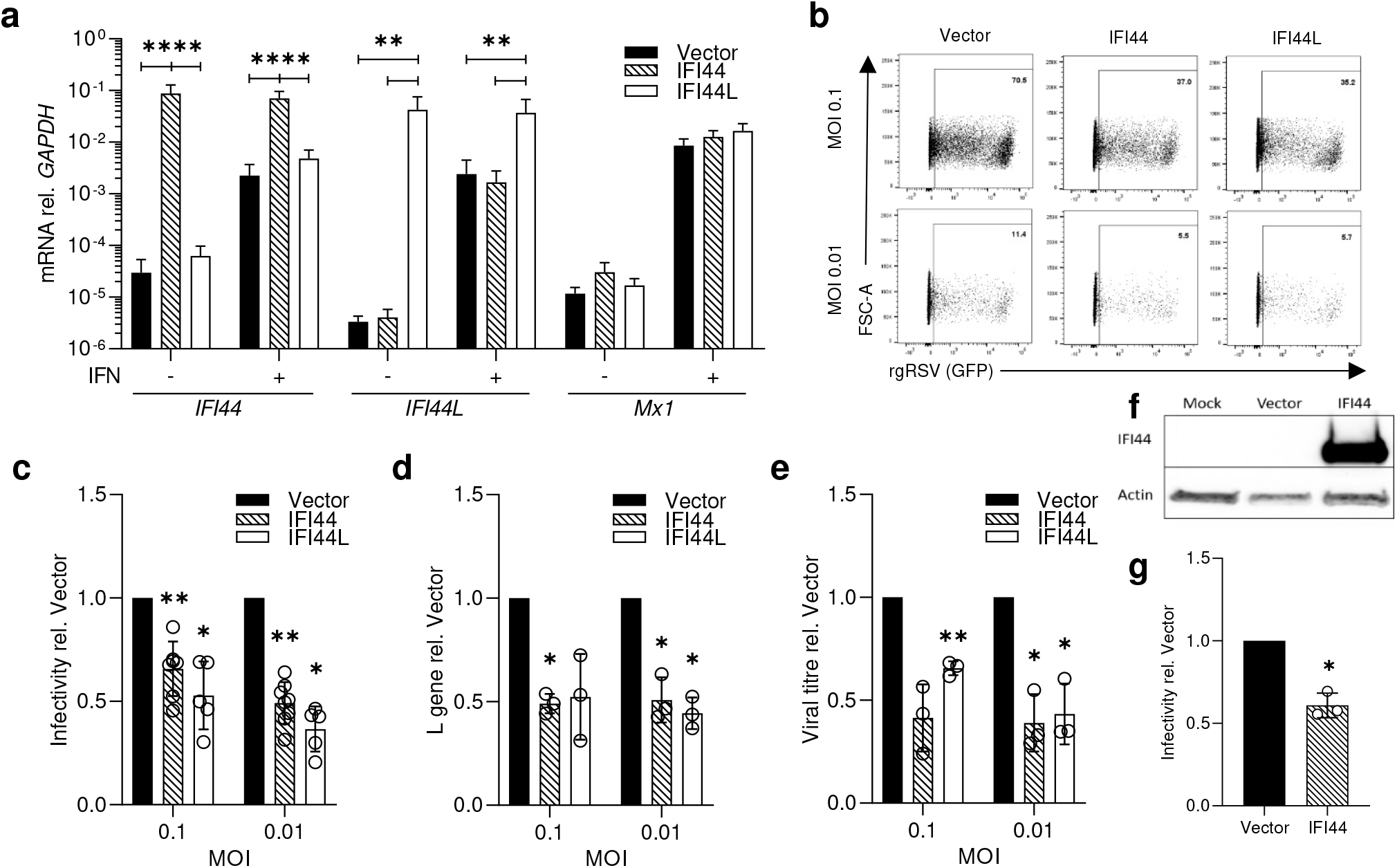
Overexpression of IFI44 or IFI44L reduces RSV infection. **(a)** *IFI44*, *IFI44L*, and *Mx1* mRNA levels relative to *GAPDH* in stably transduced overexpressing monoclonal A549 cells lines. Cells were cultured in normal growth medium or medium supplemented with 500 IU mL^−1^ IFNα2a for 24 hours. **(b)** Stably transduced cell lines were infected with rgRSV (m.o.i. 0.1 or 0.01) and infection assessed by flow cytometry after 24 hours. Representative experiment showing GFP^+^ population of single RFP^+^ cells. **(c)** RSV infection of stably transduced cell lines relative to the Vector control. N≥5. **(d)** RSV L gene 24 hours post-infection with wild-type RSV A2 relative to Vector control. **(e)** Viral titre (WT RSV A2) relative to Vector control in cell supernatant at 48 hpi by plaque assay. **(f)** A549 cells were transduced with either FLUC (Vector) or IFI44 lentivirus and expression of IFI44 detected by Western blotting after 48 hours. **(g)** Transduced A549 cells were infected with rgRSV (m.o.i. 0.8) after 24 hours, and infectivity of transduced (RFP^+^) cells assessed 24 hours after infection. N=3. Individual points represent the result of an independent experiment. Bars show the mean +/− SEM. * represents significance relative to cells transduced with empty vector, assessed by ANOVA (**a**) or ratio paired t-test (**c-e**). Analysis was done prior to data transformation. * P < 0.05, ** P < 0.01, *** P < 0.001, **** P < 0.0001.

To monitor the impact on infection, cells were infected with m.o.i. 0.1 or 0.01 recombinant GFP-expressing RSV (rgRSV), and the percentage of RFP^+^ single cells that were GFP^+^ was quantified after 24 hours. After 24 hours of rgRSV infection, cell lines expressing either IFI44 or IFI44L showed a significant reduction in the percentage of infected cells relative to cells stably transduced with empty vector (P < 0.05, Fig. 3c). To confirm the impact on RSV infection we examined the impact on wild-type RSV A2 infection in these stable cell lines. We observed a significant reduction in viral RNA 24 hours after infection (m.o.i. 0.01) (P < 0.05, Fig. 3d). To observe impact on virus progeny production, we measured the viral titre 48 hours after infection. The recoverable titre of RSV A2 virus was significantly reduced in cells expressing either IFI44 or IFI44L (P < 0.05, Fig. 3e).

In order to demonstrate that the impact of IFI44 expression is not a result of clonal differences, we transduced polyclonal A549 cells with lentivirus expressing either Firefly luciferase (FLUC) or IFI44 and subsequently infected the cells with rgRSV after 24 hours. Expression of IFI44 protein was confirmed (Fig. 3f) and significant restriction in rgRSV infection (P < 0.05, Fig. 3g) was observed similar to that seen in the stably transduced cell line. Overall these data suggest that both IFI44 and IFI44L are able to restrict RSV infection.

### Knockout of *IFI44* results in elevated RSV infection *in vitro*

To further examine the role of IFI44 or IFI44L on RSV infection we used a pool of endoribonuclease-prepared siRNA (esiRNA) targeting *IFI44* to knockdown expression. These esiRNAs reduced levels of *IFI44* mRNA by only 48% (P < 0.01) and also reduced *IFI44L* mRNA levels by 30%, although this was not statistically significant (Fig. 4a). However, esiRNA knockdown was sufficient to cause an over 2-fold increase in the levels of viral RNA in esiRNA-IFI44 transfected A549 cells infected with RSV A2 (m.o.i. 0.1) relative to cells transfected with a non-targeting control (P < 0.01, Fig. 4b).

**Figure 4.**
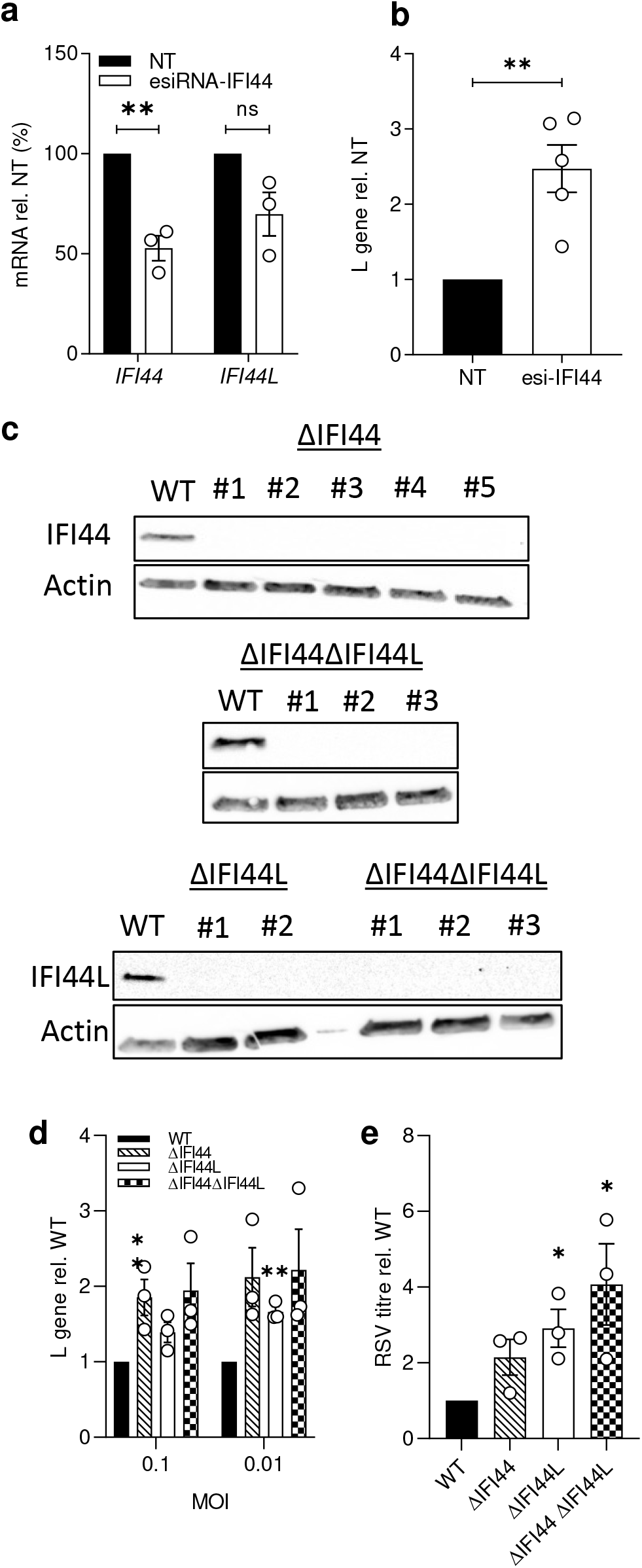
Loss of IFI44 expression enhances RSV infection *in vitro*. **(a)** *IFI44* and *IFI44L* mRNA expression after transfection with 50 nM esiRNA-IFI44 or a non-targeting esiRNA control (NT). 24 hours after transfection cells were treated with 500 IU mL^−1^ IFNα2a for 16 hours prior to infection with RSV A2 (m.o.i. 0.1) for 48 hours. N=3. **(b)** L gene copy number assessed 48 hours after infection and shown relative to the NT control. N=5. **(c)** A549 monoclonal knockout cell lines were generated through CRISPR-Cas9 gene editing. IFI44 or IFI44L protein in wild-type cells treated with IFNα2a or monoclonal CRISPR-Cas9 edited A549 cells. **(d)** Cells were treated with 500 IU mL^−1^ IFNα2a 16 hours prior to infection with RSV A2 (m.o.i. 0.1 or 0.01) for 24 hours. RSV L gene copy number shown relative to the WT control. N=3. **(e)** Knockout cell lines were pre-treated with IFNα2a as for **c** and subsequently infected with RSV A2 (m.o.i. 0.1) for 48 hours. Viral titre in cell supernatant relative to the WT control. N=3. Points represent a single independent experiment with a bar at the mean +/− SEM. Significance relative to the WT or NT controls by ratio paired T test. Analysis was done prior to data transformation. * P < 0.05, ** P < 0.01. ns = not significant.

Due to the low efficiency of the knockdown and potential unintended impact of siRNA on *IFI44L* expression we developed CRISPR-Cas9 edited IFI44 and IFI44L deficient cell lines, this also enabled us to investigate the impact of double knockouts. A549 cells were transfected with two guide RNA (gRNA) constructs encoded *in cis* with the *Streptococcus pyogenes* Cas9 enzyme and a GFP marker. gRNA sequences targeting regions in exons 4 and 5 of *IFI44* and exons 3 and 4 of *IFI44L* were used. Following transfection, the top 2% GFP^+^ cells were sorted as single cells. Gene editing was confirmed in clonal populations through PCR of the targeted region (data not shown) and loss of either IFI44 or IFI44L protein expression after treating with IFNα (Fig. 4c). Five IFI44 deficient (ΔIFI44), two IFI44L deficient (ΔIFI44L), and three clonal populations deficient in both IFI44 and IFI44L (ΔIFI44ΔIFI44L) were isolated. Selection of clones was based on sequencing. The ΔIFI44 clone selected for further study had mutations resulting in a frameshift and stop codon formation in each detectable allele. The ΔIFI44L clone selected had a large deletion in one allele and disruption of an exon-intron boundary.

The selected clones were treated with 500 IU ml^−1^ IFNα2α and subsequently infected with RSV A2 for 24 hours (Fig. 4d). There were significantly increased levels of viral RNA in the ΔIFI44 clone at m.o.i. 0.1 (P < 0.01), although we noted a non-significant increase in infection in the ΔIFI44ΔIFI44L cells (P = 0.079). The loss of IFI44L expression was only associated with a significant increase in viral RNA at m.o.i. 0.01 (P < 0.01). Clones were infected with RSV A2 (m.o.i. 0.1) for 48 hours and viral titre assessed in the culture medium, there was with a >2-fold increase in viral titre (Fig. 4e) in each clone. Knockout of both *IFI44* and *IFI44L* resulted in a 4-fold increase in viral titre (Fig. 4e, P < 0.05).

### IFI44 and IFI44L reduce cellular proliferation

Previous studies have described an effect of IFI44 or IFI44L on cell proliferation (13, 18). We investigated the impact of these factors on proliferation by assaying growth of selected knockout clones. Knocking out either or both genes was associated with increased proliferation, using a colorimetric assay measuring cellular metabolic activity (P < 0.05, Fig. 5a). When viable cell numbers were quantified manually by Trypan blue exclusion only the *IFI44* KO was associated with a significant increase in cell number (P < 0.05, Fig. 5b). Overexpression of either gene led to a significant reduction in proliferation after 24 hours quantified by either method (P < 0.05, Fig. 5c-d). Cells perfused with CellTrace Violet dye were allowed to proliferate for 72 hours prior to analysis by flow cytometry to assess cell division. We noted that both IFI44 and IFI44L stably transduced cell lines had an increased mean fluorescence intensity (P < 0.05, Fig. 5f) suggesting reduced dye dilution and a reduced rate of cell division. The overexpression of IFI44 or IFI44L was not associated with any increase in cytotoxicity, further suggesting that the observed reduction in viable cell number and proliferation is not a result of increased cell death (Fig. 5g).

**Figure 5.**
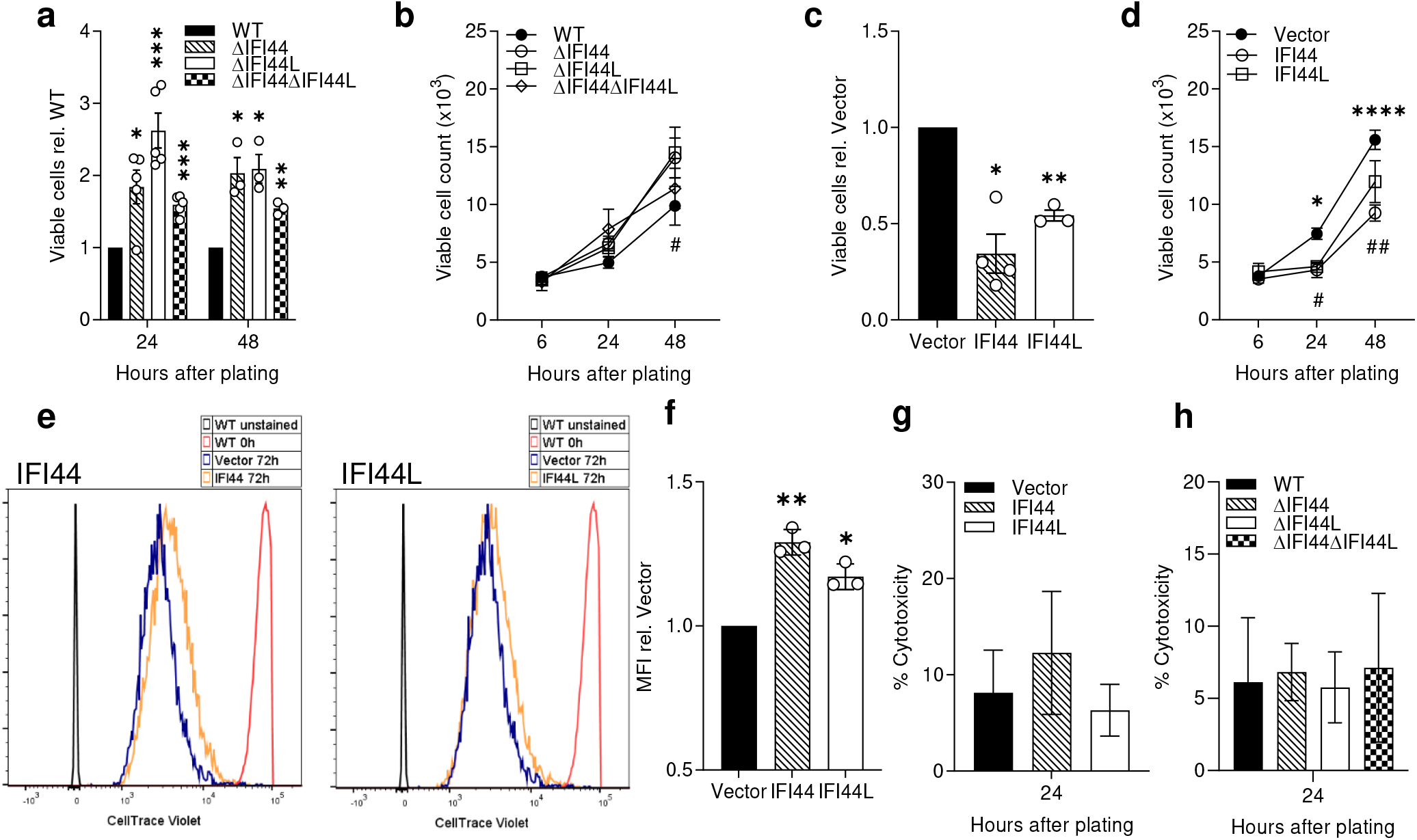
IFI44 and IFI44L are anti-proliferative. A549 monoclonal knockout cell lines were seeded at equal densities and viable cell number quantified after 6-48 hours using a colorimetric metabolic activity assay **(a)** or by trypan blue exclusion (# P < 0.05 between ΔIFI44L and vector). **(b)**. A549 stably transduced monoclonal cell lines expressing IFI44, IFI44L, or transduced with empty vector were seeded at equal densities and viable cell number quantified 24 hours later by colorimetric metabolic activity assay **(c)** or after 6-48h by trypan blue exclusion **(d)** (* P <0.05, **** P < 0.0001 between IFI44 and vector, # P <0.05, ## P <0.01 between IFI44L and vector) N≥3. **(e)** Representative histograms of stably transduced A549 cell lines treated with CellTrace Violet and allowed to proliferate for 72 hours before analysis by flow cytometry. WT cells were stained or treated with vehicle only and immediately fixed prior to analysis for positive (red) and negative (grey) controls. **(f)** Mean fluorescence intensity (MFI) of cell trace violet quantified relative to Vector control over three independent repeats. Cytotoxicity was assessed by LDH release assay 24-48 hours after plating in overexpression **(g)** or knockout cells **(h)**. N=3. Significance to WT or Vector controls assessed by ratio paired T test prior to data transformation. Points represent a single independent experiment with a bar at the mean +/− SEM. * P < 0.05, ** < P <0.01, *** P < 0.001, **** P < 0.0001 in panels a, c and f between indicated bar and vector.

### IFI44 reduces RSV polymerase activity

To investigate where in the viral life cycle IFI44 and IFI44L have an impact we analysed infection at an acute time point (8h) where viral positive cells should only be newly infected cells and not the result of cell-cell virus spread. IFI44 or IFI44L expression reduced the percentage of infected cells by 44% (P < 0.01) and 34% (P = 0.11) respectively (Fig. 6a-b) relative to vector control cells, suggesting both proteins are impacting a stage of the viral life cycle prior to new virion release. Using a cold-bind infection assay (RSV A2, m.o.i. 2), where the virus is able to bind the cell surface but is not internalised, we saw no significant difference in levels of viral RNA between cell lines expressing either FLUC (Vector), IFI44, or IFI44L (Fig. 6c). To bypass cell entry and to assess whether IFI44 or IFI44L restrict RSV genome replication or transcription we transfected stably transduced clonal 293T cells with an RSV minigenome system. IFI44 expression reduced minigenome activity by 44% (P < 0.05), suggesting reduced RSV polymerase activity (Fig. 6d). The stably transduced 293T line expressing IFI44L also reduced minigenome activity by 45% although this was not statistically significant (P = 0.085).

**Figure 6.**
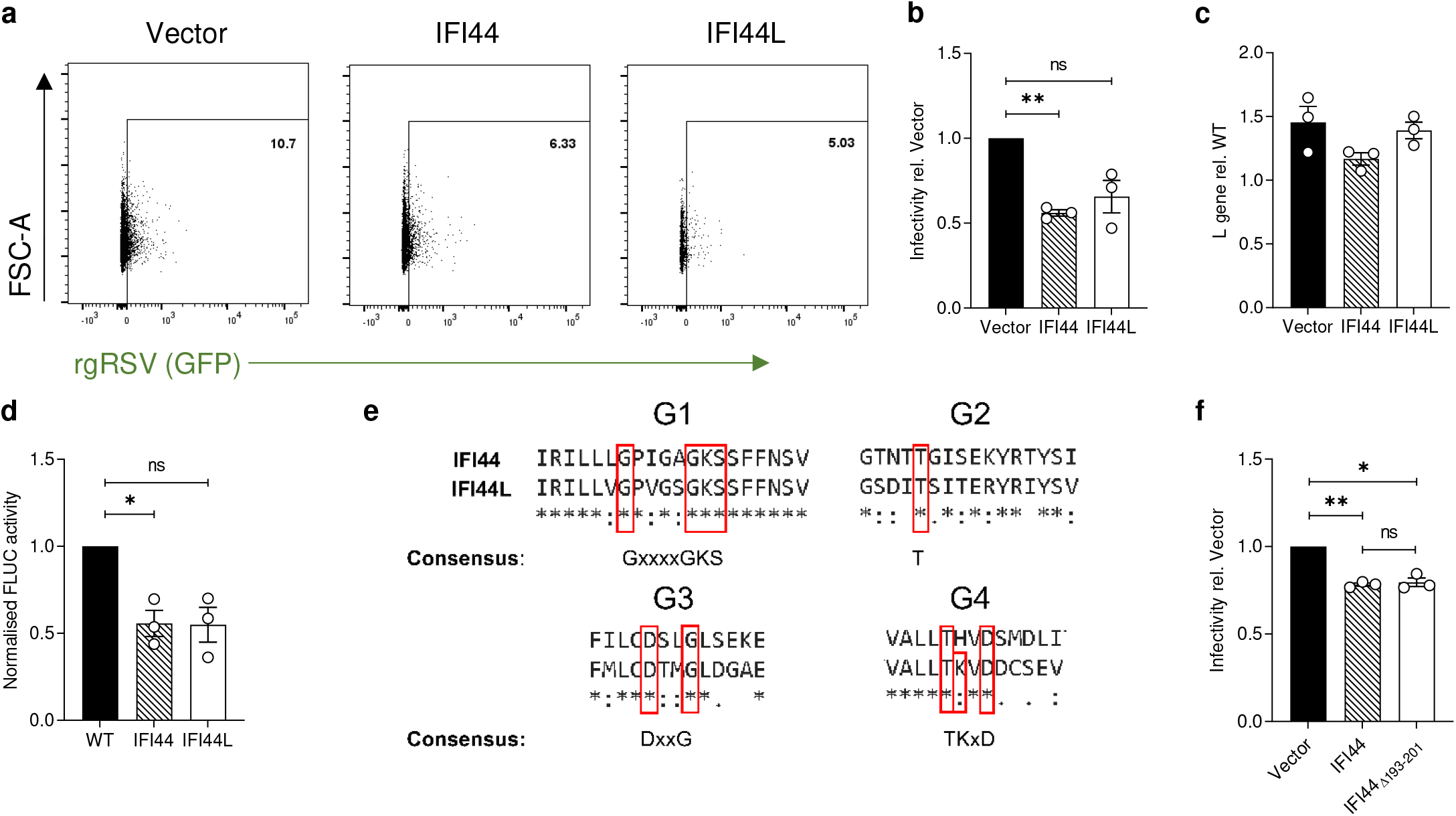
IFI44 and IFI44L reduce RSV polymerase activity and restrict infection independent of GTP-binding. **(a)** Stably transduced A549 cells were infected with rgRSV (m.o.i. 0.1) for 8 hours and infectivity assessed by measuring the percentage of RFP^+^ single cells that were GFP^+^. Representative dot plot from a single independent experiment. Population shown was gated for single RFP^+^ cells. **(b)** Quantification of rgRSV infection relative to Vector transduced control as described in **a**. N=3. **(c)** Stably transduced A549 cell lines were incubated with RSV A2 (m.o.i. 2) for 1 hour at 4 °C then harvested for analysis of RSV L gene copy number. N=3. **(d)** Stably transduced 293T cell lines were transfected with pSV-β-Gal, pCAGGS-T7, the pGEM3-Gaussia/Firefly minigenome and plasmids encoding RSV M2-1, P, L, and N. 24 hours later FLUC activity was assessed and normalised to the negative control and β-Galactosidase expression levels. Normalised FLUC activity shown relative to polyclonal parental 293T cells (WT). N=3. **(e)** ClustalW multiple sequence alignment of human IFI44 and IFI44L. Four sections were selected to show the residues predicted to be essential for GTP binding and GTPase function (highlighted by a red box). Consensus sequence for each G1-G4 region shown below. **(f)** A549 cells were transduced with 2×10^5^ TU lentivirus expressing either WT IFI44, IFI44Δ193-201 (G1 region deleted), or FLUC (Vector). 24 hours later cells were infected with rgRSV (m.o.i. 0.8) and infectivity assessed as for **a** 24 hours after infection. N=3. Significance to Vector transduced or WT controls assessed by ratio paired T test prior to data transformation. Points represent a single independent experiment with a bar at the mean +/− SEM * P < 0.05, ** < P <0.01. ns = not significant.

### A predicted GDP/GTP-binding site in IFI44 is dispensable for antiviral activity

IFI44 and IFI44L both contain predicted GTPase regions. Five motifs, G1-G5, are involved in GTP hydrolysis and the required nucleotide and cofactor interactions (32). Both proteins contain a putative G1 region (GXXXXGKS), initially identified in IFI44 by Hallen *et al* (13), that is predicted to bind the β-phosphate of GDP or GTP (32). The terminal serine residue also contributes to Mg^2+^ binding. The G1-G4 regions of human IFI44 and IFI44L were recently annotated by McDowell *et al* (Fig. 6e) (33). The G3 DxxG motif in IFI44 and IFI44L is immediately followed by a hydrophobic leucine residue found in other dynamin-like GTPases such as GBP1, instead of a catalytic glutamine residue seen in ras-like GTPases. IFI44L also contains a G4 motif which mediates binding to the guanine base of GDP or GTP. However, IFI44 does not contain a complete G4 motif. G5 residues are not easily recognised as they are not as well conserved as G1-4 motifs.

A549 WT cells were transduced with a lentivirus expressing FLUC (Vector), IFI44 WT, or IFI44_Δ193-201_ with the G1 region deleted. Mutation of this region in another IFN-inducible GTPase, GBP1, has been shown to reduce nucleotide binding around up to 50-fold, having a greater impact than mutation of G2-G4 regions (34). A comparable reduction in rgRSV infectivity was seen in cells transduced with either IFI44_Δ193-201_ or IF44 WT (Fig. 6f), suggesting this region is not required for its antiviral function.

### Disease severity is altered in an *Ifi44*^*−/−*^/*Ifi44L*^*−/−*^ mouse model of RSV infection

Having demonstrated that IFI44 and IFI44L were able to impact RSV infection *in vitro* we then investigated the effect of the absence of *Ifi44* and *Ifi44L* in an *in vivo* mouse model of RSV infection. The wild-type C57BL/6N used in this study are Ifi44L^−/−^, presumably as a result of gene loss over colony in-breeding, so the comparison was between IFI44^−/−^/IFI44L^−/−^ and IFI44^+/+^/IFI44L^−/−^ (WT). Age-matched *Ifi44^−/−^* and WT mice were infected intranasally with 10^5^ PFU RSV A2 and monitored for weight change over a 7-day infection. Animals were sacrificed at day 4 and 7 after infection. There was greater weight loss in *Ifi44*^*−/−*^ mice than wild-type controls from days 5-7 after infection (P < 0.01, Fig. 7a). There was no difference in total cell numbers isolated from the bronchoalveolar lavage fluid or from whole lung tissue (Fig. 7b-c). Levels of viral RNA in lung tissue was also significantly higher in *Ifi44*^*−/−*^ mice at day 4 (P < 0.01) but not at day 7 (Fig. 7d).

**Figure 7.**
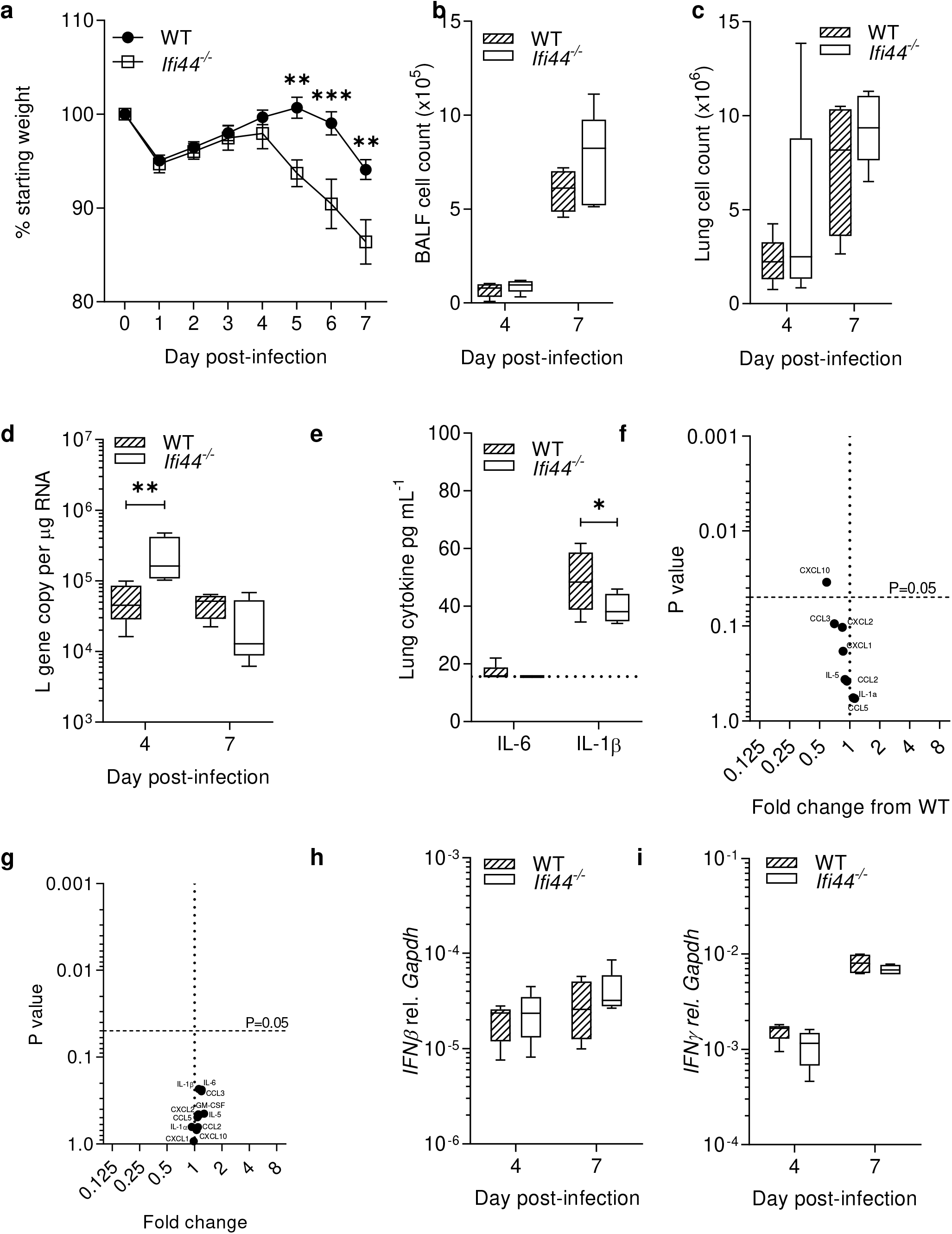
RSV infection severity is enhanced in a *Ifi44*^*−/−*^ mouse model. **(a)** Wildtype or *Ifi44*^*−/−*^ C57BL/6N mice were infected intranasally with 1 × 10^5^ pfu RSV A2. Weight loss was monitored for 7 days. **(b)** BALF and **(c)** lung cell counts. **(d)** Viral load assessed by RSV L gene qPCR. **(e)** IL-6 and IL-1β levels at day 4 post-infection measured by ELISA. Volcano plot of inflammatory cytokines at day 4 **(f)** and day 7 **(g)** by multiplex ELISA. **(h)** *Ifnβ* and **(i)** *Ifnγ* mRNA relative to *Gapdh* (2^-ΔCt). Box plots show a line at the median and box edges from the 25^th^ to 75^th^ percentiles, with whiskers from the 5^th^ to 95^th^ percentiles (Prism). N≥4 animals at each time point. Two independent experiments. Significance by ANOVA (**a**-**e**, **h**-**i**) or Student’s T test (**f-g**). * P < 0.05, ** 0.01, *** P < 0.001.

Levels of inflammatory cytokines and immunomodulatory factors were analysed either by ELISA or Luminex. Here we noted that most measured analytes (IL-6, CCL3, CXCL2, CXCL1, CCL2, IL-5, IL-1α, and CCL5) were not significantly different between the WT and *Ifi44*^*−/−*^ groups (Fig. 7e-f). CXCL10 (P < 0.05, Fig. 7f) and IL-1β (P < 0.05, Fig. 7e) were both significantly but modestly reduced in the KO animals on day 4 after infection. There was no difference in any measured cytokine on day 7 after infection (Fig. 7g) To determine whether the lack of IFI44 modulated IFN responses to RSV infection both *Ifn-β* and *Ifn-γ* expression was assessed by qPCR. Both wild-type and *Ifi44*^*−/−*^ groups demonstrated similar levels of *Ifn-β* (Fig. 7h) and *Ifn-γ* (Fig. 7i) mRNA.

## Discussion

The data presented here explore how the ISGs IFI44 and IFI44L modulate viral infection. We demonstrate that IFI44 and IFI44L restrict RSV infection and reduce RSV genome replication or transcription. We also show, for the first time, that RSV infection is enhanced in an *Ifi44*^*−/−*^/*Ifi44L*^*−/−*^ knockout mouse model. Infectivity was reduced by IFI44 expression at just 8 hours after infection, suggesting restriction of infection occurs before the exit of new virions. Virus attachment was unaffected by either IFI44 or IFI44L expression, this was expected as these proteins are both predicted to be internally expressed (12, 13). Using an RSV minigenome assay, we found that IFI44 expression significantly reduced RSV polymerase activity. However, we cannot say whether this is an impact specifically on the replication or transcription of the viral genome. We also observed that both IFI44 proteins decreased the rate of cellular proliferation. Reduced proliferation is a common feature of the IFN response, mediated by canonical ISGs such as protein kinase R (PKR) (35) and the IFN-induced tetratricopeptide repeat (IFIT) family (36). Whether the anti-proliferative function of the IFI44 proteins is a causative mechanism of their antiviral activity is not clear, because cell cycle arrest may increase the availability of cellular machinery required for replication and virus assembly (37). For example, the RSV matrix protein (M) has previously been shown to induce cell cycle arrest by inducing p53/p21 expression in alveolar epithelial cells, enhancing infection (38, 39). The two genes have a high degree of homology and whether they have distinct mechanisms or are redundant is unclear at this time.

Our *in vitro* studies have some limitations, primarily the use of clonal cell lines to assay infection and proliferation. We note that transduction of polyclonal parental A549 cells with IFI44 is capable of restricting RSV infection similarly to the stably transduced cells, and that previous studies have observed similar impacts of IFI44 on proliferation (13, 18) or restriction of RSV (8). However, it is possible that clone-specific differences have some impact on either RSV infection or cell viability and these data should be interpreted with this limitation in mind. In the knockout cell line, there was a slight difference in the effect on viral RNA and infectious virus recovered, this may reflect differences in the two assays or a difference in where the ISG affect replication or packaging.

This is the first study to describe viral infection in *Ifi44*^*−/−*^/*Ifi44L*^*−/−*^ mice. *Ifi44*^*−/−*^/*Ifi44L*^*−/−*^ mice were markedly more susceptible to RSV infection than WT mice, exhibiting increased weight loss and elevated viral load. The genome sequence of C57BL/6N mice reveals a deletion in *Ifi44l* predicted to ablate expression. We were unable to detect transcription of this gene in these mice during RSV infection whereas this was readily detectable in the lungs of RSV-infected BALB/c mice. We observed that *Ifi44*^*−/−*^/*Ifi44L*^*−/−*^ mice infected with RSV had higher levels of viral RNA present in their lungs at the peak of infection, along with decreased expression of the pro-inflammatory factor IL-1β. A previous study has noted that in adult mice, blockade of IL-1β prior to RSV infection, results in elevated viral load (40). Decreased production of this key cytokine along with increased viral replication, due to changes in cellular proliferation and metabolism driven by the loss of IFI44, may go some way to explaining these observations. We saw no change in TNF, but this reflects our recent findings that TNF is only associated with early weight loss after RSV infection, but not later time points (41).

Our data showing an antiviral role for IFI44 and IFI44L matches our previous study using a lentivirus screen (8) and a broader screen by another group using the same lentivirus panel (14). However, the data presented here is somewhat at odds with recently published studies investigating the impact of IFI44 (42) and IFI44L (43) on virus in vitro. In the published studies reducing IFI44 or IFI44L in vitro using siRNA led to increased viral recovery, which was hypothesized to be linked to decreased ISG expression. We did not see an effect of IFI44 or IFI44L over-expression on the expression of the ISG Mx1. One possible difference between study designs is the role of cell proliferation. We observed that IFI44 and IFI44L had a significant anti-proliferative effect and to normalise we counted cell numbers in parallel wells prior to infections and altered the viral inoculum to ensure equivalent MOI were used. The parallels between the in vivo and in vitro phenotype give us confidence that the genes can have an antiviral function.

One question of interest is about the redundancy of ISGs in the control of infection. Using knockout mouse models, an increase in RSV disease severity has been seen for a variety of antiviral ISGs such as Ifitm3 (44), Ifitm1 (5), and Irf7 (8). It is curious that in these *in vitro* and *in vivo* models, single gene loss can result in the loss of viral control, when the host network of ISGs consists of potentially over 1000 genes. Whilst some associations between individual ISGs and disease severity have been observed in humans – for example IFITM3 (45, 46), more often primary immunodeficiencies caused by mutations in the interferon sensing and signalling pathways such as STAT1 and TLR3 display incomplete penetrance and only specific susceptibilities (47). Potentially the use of large volumes relative to lung size and high doses of virus to ensure infection in the mouse model stresses the system and therefore the role of individual genes becomes more apparent. It was of note that we observed a significant reduction in IFI44 and IFI44L expression associated with infant patients requiring PICU admission, when tested in isolation, but in the global comparison, there were other genes that were more significantly different, suggesting there may be a more coordinated effect that influences disease severity. It should also be noted that the children requiring intensive care were significantly younger, which may impact gene expression. This may be the result of reduced IFN activity as a whole – this has been seen in another cohort of children with RSV infection (48) where those children with severe infection had lower global expression of ISG. This suggests that ISGs work in concert to control infection, targeting different aspects of the viral life cycle, with some factors such as IFITM1 controlling entry (5) and others restricting replication within the cell. Further work is required to understand whether lower expression levels during human infection predispose to severe RSV and might explain some of the heterogeneity in disease severity in children. Several expression quantitative trait loci (eQTL) are described for these genes (49), including one located within the *IFI44L* gene, rs273259, that was the top-hit in a genome-wide association study of adverse responses to a live virus vaccine (MMR) (50). Cells with the rs273259 risk allele expressed *IFI44* at lower levels, and *IFI44L* with altered splice isoform levels (50). These findings establish a precedent for polymorphisms affecting *IFI44* and *IFI44L* expression to influence predisposition to a severe response to viral infection.

Our study demonstrates that IFI44 and IFI44L play a role in the control of RSV infection. They work intracellularly, reducing the ability of the virus to replicate. Since the proteins are anti-proliferative, this may be part of the mechanism. Understanding how they reduce viral replication may provide future avenues for therapeutic interventions.

## Acknowledgements

Marcus Dorner and Jessie Skelton for providing lentiviral constructs for gene expression. DCB studentship was funded by The Wellcome Trust (109056/Z/15/A).

## Contributions

DCB: Conceptualization, Investigation, Methodology, Writing – original draft
DH-C: Formal analysis
SC: Resources, Investigation
CB: Investigation
IB: Resources
MK: Formal analysis, Supervision
JH: Resources
ML: Supervision, Funding Acquisition
JFE: Resources
PK: Supervision, Writing-Review and editing
JST: Conceptualization, Funding Acquisition, Writing – original draft.

